# Linking lysosomal enzyme targeting genes and energy metabolism with altered gray matter volume in children with persistent stuttering

**DOI:** 10.1101/848796

**Authors:** Ho Ming Chow, Emily O. Garnett, Hua Li, Andrew Etchell, Jorge Sepulcre, Dennis Drayna, Diane C. Chugani, Soo-Eun Chang

## Abstract

Developmental stuttering is a childhood onset neurodevelopmental disorder with an unclear etiology. Subtle changes in brain structure and function are present in both children and adults who stutter. It is a highly heritable disorder, and up to 12-20% of stuttering cases may carry a mutation in one of four genes involved in mannose-6-phosphate mediated protein intracellular trafficking. To better understand the relationship between genetic factors and brain structural changes, we used gene expression data from the Allen Institute for Brain Science (AIBS) and voxel-based morphometry (VBM) to investigate the spatial correspondence between gene expression patterns and differences in gray matter volume (GMV) between children with persistent stuttering (n=26, 87 scans) and their fluent peers (n=44, 139 scans). We found that expression patterns of two stuttering-related genes (*GNPTG* and *NAGPA*) in the brain exhibit a strong positive spatial correlation with the magnitude of GMV differences between groups. Further gene set enrichment analyses revealed that genes whose expression was highly correlated with the GMV differences were enriched for glycolysis and oxidative metabolism in mitochondria. Although the results are correlational and cannot inform us about underlying casual mechanisms, our results suggest a possibility that regions with high expression level of genes associated with stuttering may be particularly vulnerable to the effect of alterations in these genes. This effect may be further exacerbated by the relatively high energy utilization in those brain during the period of a sharp increase in brain energy utilization, which coincides with a period of rapid language development and the onset of stuttering during childhood.

## Introduction

Fluid, effortless speech production forms the basis for communication and is considered a fundamental human ability. Stuttering significantly disrupts fluent speech production, often leading to negative psychosocial and economic consequences throughout life (Blumgart, Tran, & Craig, 2010; Craig, Blumgart, & Tran, 2009; Yaruss, 2010). Developmental stuttering typically has an onset in early childhood, affecting more than 5% of preschool-age children and persisting in about 1% of adults (Craig, Hancock, Tran, Craig, & Peters, 2002; Månsson, 2000; Yairi & Ambrose, 2013). Persistent stuttering is highly heritable, with estimations of genetic contribution exceeding 80% in some studies (Dworzynski, Remington, Rijsdijk, Howell, & Plomin, 2007; Fagnani, Fibiger, Skytthe, & Hjelmborg, 2011; Ooki, 2005; Rautakoski, Hannus, Simberg, Sandnabba, & Santtila, 2012; van Beijsterveldt, Felsenfeld, & Boomsma, 2010). Genes causative of persistent stuttering have begun to be identified (C. Kang et al., 2010; Raza et al., 2015). To date, four such genes, designated *GNPTG*, *GNPTAB* and *NAGPA* and *AP4E1* have been found, and they may cumulatively account for up to 12-20% of unrelated individuals with persistent stuttering (see Frigerio-Domingues & Drayna, 2017 for a comprehensive review). This group of genes is known to play a role in lysosomal enzyme trafficking. *GNPTG*, *GNPTAB* and *NAGPA* are involved in marking lysosomal hydrolases and several nonlysosomal proteins with a mannose-6-phosphate (M6P) tag that is important for intracellular trafficking (Barnes et al., 2011). Homozygous mutations in *GNPTG* and *GNPTAB* genes are known to cause the rare inherited lysosomal storage disorders Mucolipidosis Types II and III (Kornfeld, 2001), which affect many parts of the body including the brain. However, in most of the cases, people who stutter only carry heterozygous mutations in these genes, and do not have the signs or symptoms typically seen in Mucolipidosis types II and III. *AP4E1* is a member of a family of adaptor proteins that are involved in vesicle formation and sorting member proteins for transporting lysosomal enzymes from the trans Golgi network to late endosomes and lysosomes. Mutations in *AP4E1* have been associated with hereditary spastic paraplegia and cerebral palsy (Abou Jamra et al., 2011; Kong et al., 2013). Why mutations in these genes specifically affect the ability to produce fluent speech but not other cognitive or neurologic functions remains unknown. However, neuroimaging studies have shown that persistent stuttering is associated with subtle functional and anatomical anomalies (Chow & Chang, 2017; Garnett et al., 2018; Neef, Anwander, & Friederici, 2015). How genetic factors relate to brain anomalies is not yet clear. To pursue this question, we examined the differences in spatial patterns of gray matter volume (GMV) in children with persistent stuttering (pCWS) with the regional expression of the four genes thus far associated with stuttering using data provided by the Allen Institute for Brain Science (AIBS; http://www.brain-map.org/). This approach has been used to reveal gene-brain relationships in a number of recent studies. A seminal study was published in 2015, in which the authors used gene expression data and resting-state functional magnetic resonance imaging (fMRI) to identify 136 genes associated with intrinsic functional networks in the brain (Richiardi et al., 2015). In another study, Ortiz-Terán et al. used a similar method to demonstrate that the neural reorganization in blind children measured by resting-state functional connectivity is associated with a set of known neuroplasticity-related genes (Ortiz-Terán et al., 2017). Moreover, this approach was also employed to study neuropsychiatric disorders. For example, McColgan et al. identified genes associated with Huntington’s disease by comparing regional white matter loss and gene expression in patients with the disorder (McColgan et al., 2017).

Although we do not know the mutation status of our participants with stuttering, high heritability of the disorder suggests that they are likely to carry a known or yet to be discovered gene mutation. While the genetic causes of stuttering are likely to be heterogeneous, it is probable that their effects at the neuroanatomical level could be similar because the disorder affects speech production specifically. Moreover, in support of the previous argument, neuroimaging studies of people who stutter with unknown genetic status have shown some consistent results. In particular, different groups of researchers have independently demonstrated that the anisotropic diffusivity in the corpus callosum and the superior longitudinal fasciculus are decreased in people who stutter compared with matched controls (Neef et al., 2015). Therefore, a core presumption of this study is that the expression patterns of the known genes associated with stuttering, to a certain extent, reflect the magnitude of anatomical anomalies of the disorder. A similar presumption was made in a previous study showing that the expression of risk genes for schizophrenia were positively correlated with the anatomical disconnectivity defined by diffusion tensor imaging (DTI) tractography in patients with schizophrenia, but not in patients with bipolar disorder (Romme, de Reus, Ophoff, Kahn, & van den Heuvel, 2017).

In this study, we hypothesized that the expression of the four stuttering-related genes would be associated with the GMV differences in pCWS. Moreover, genes whose expression is highly associated with the GMV differences were used to explore the potential the biological processes and pathways involved in stuttering, using gene set enrichment analysis.

## Materials and Methods

### Participants

Participants were primarily recruited from the East Lansing, Michigan area and surrounding 50-mile radius, as part of an on-going longitudinal study conducted at Michigan State University (MSU). Recruitment activities included contacting and sending study flyers to area physicians’ offices, childcare facilities, public schools, and outpatient speech clinics, and advertising in parent magazines and in social media. Relevant to the current investigation, a total of 226 children were contacted and screened. Of those, 128 were eligible and willing to participate in the study. Thirty-three of 128 did not participate in the MRI for variable reasons (e.g., uneasiness in the MRI setting and anxiety detected during mock scanner training), while 50 children who stutter (CWS; 20 girls and 30 boys) and 45 controls (23 girls and 22 boys) were scanned. Each subject participated in 1 to 4 longitudinal visits, with an inter-visit interval of approximately 12 months. The mean ages of CWS and controls at the first visit were 5.55 (SD=2.02) and 5.99 (SD=2.00) years, ranging between 3 to 10 years. Participants were monolingual English speakers. Children in the two groups did not differ in chronological age, sex, handedness, or socioeconomic status (SES) based on mother’s education level. All children exhibited normal speech and language development except for the presence of stuttering in the stuttering cohort as confirmed through a battery of standardized assessments, including Wechsler Preschool and Primary Scale of Intelligence Third Edition for children 2:6-7:3 (Wechsler, 2002), Wechsler Abbreviated Scale of Intelligence for children 7 and up (Wechsler, 1999), Peabody Picture Vocabulary Test (PPVT-3) for receptive vocabulary ability (Dunn & Dunn, 2007), Expressive Vocabulary Test (EVT-2; Williams, 2007) and Goldman-Fristoe Test of Articulation (GFTA-2; Goldman, 2000). The results of these tests are listed in Table 1. For study inclusion the participants had to score above −2 standard deviations of the norm on all standardized tests. None of the subjects had any concomitant developmental disorders (e.g., dyslexia, ADHD), with the exception of stuttering in CWS, and none were taking any medication affecting the central nervous system. CWS with assessments from 2 visits or more were further categorized as recovered or persistent based on their stuttering severity rating. The Stuttering Severity Instrument (SSI-4) was used to examine frequency and duration of disfluencies occurring in the speech sample, as well as any physical concomitants associated with stuttering. These were incorporated into a composite stuttering severity rating (Riley & Bakker, 2009). A child was considered recovered if the SSI-4 score was 10 or below at the second visit or thereafter. A child was categorized persistent if the composite SSI-4 score was higher than 10 at the second visit or thereafter. Both clinician and parent reports were required to be consistent with stuttering severity assessments in determining whether a child had recovered or was persistent. Three CWS were excluded because they were assessed only once and, therefore their final diagnoses could not be determined. Another CWS was excluded due to incidental findings in the structural scan. Nine scans from seven subjects were excluded due to excessive head movements. The final analysis included 87 scans from 26 children with persistent stuttering (pCWS; 8 girls and 18 boys; mean age at the first visit=6.5 years; SD=1.9), 61 scans from 17 children recovered from stuttering (rCWS; 8 girls and 9 boys; mean age at the first visit= 5.4 years; SD=1.9), and 139 scans from 44 controls (23 girls and 21 boys; mean age at the first visit=6.5 years; SD=2.0). Both persistent and recovered groups did not differ from controls in chronological age, sex, handedness, or SES. Because *GNPTAG*, *NAGPA*, *GNPTG* and AP4E1 are associated with persistent stuttering, only scans collected from children with persistent stuttering and controls were included in the primary analysis. All research procedures were approved by the Michigan State University Institutional Review Board, which follows the ethical standards described in the Belmont Report and complies with the requirements of the Federalwide Assurance for the Protection of Human Subjects regulated by the United States Department of Health and Human Services. Written consents were obtained from all parents of the participating children, and assents were obtained from all children in verbal or written format depending on reading level. Children were paid a nominal remuneration, and were given small prizes (e.g., stickers) for their participation.

**Table 1.** Demographics, intelligent quotient (IQ) and language tests scores averaged across longitudinal visits for each participant

|  | Controls, n = 44 (21 boys) |  | Persistent, n = 26 (18 boys) |  | Recovered, n = 17 (9 boys) |  |
| --- | --- | --- | --- | --- | --- | --- |
|  | Mean (SD) | Range | Mean (SD) | Range | Mean (SD) | Range |
| Age at the first scan (years) | 6.5 (2.0) | 3.3 – 10.8 | 6.5 (1.9) | 3.6 – 10.3 | 5.4 (1.9) | 3.1 – 9.4 |
| IQ <sup>a</sup> | 114 (14.1) | 84 – 144 | 106 (15.5) | 81 – 138 | 106 (13.1) | 88 – 128 |
| PPVT <sup>b</sup> | 119 (12.7) | 95 – 141 | 110 (13.5) | 86 – 146 | 114 (10.3) | 93 – 131 |
| EVT <sup>c</sup> | 115 (11.8) | 93 – 142 | 106 (12.2) | 86 – 138 | 109 (9.2) | 89 – 129 |
| GFTA <sup>d</sup> | 104 (6.6) | 84 – 115 | 102 (4.2) | 92 – 110 | 106 (7.3) | 91 – 115 |
| SSI <sup>e</sup> | - | - | 21 (8.3) | 12 – 48 | 13 (2.9) | 7 – 19 |

<sup>a</sup> intelligence quotient (IQ) No significant difference between any two groups (t-tests, $p > 0.05$ )
<sup>b</sup> The Peabody Picture Vocabulary Test (PPVT). Scores significantly higher in control than persistent groups (two-sample t-tests, $p < 0.05$ ).
<sup>c</sup> The Expressive Vocabulary Test (EVT). Scores significantly higher in control than both persistent and recovered groups (two-sample t-tests, $p < 0.05$ ).
<sup>d</sup> The Goldman-Fristoe Test of Articulation (GFTA). Scores significantly higher in control than persistent groups (two-sample t-tests, $p<0.05$ ).
<sup>e</sup> Scores significantly higher in persistent and recovered groups (two-sample t-tests, $p<0.05$ ).

### Voxel-based morphometry (VBM)

Anatomical images were acquired on a GE 3T Signa scanner with an 8-channel head coil at Michigan State University. In each scan session, a whole brain 3D inversion recovery fast spoiled gradient-recalled T1-weighted images with CSF suppressed was obtained using the following parameters: time of echo = 3.8 ms, time of repetition of acquisition = 8.6 ms, time of inversion = 831 ms, repetition time of inversion = 2,332 ms, flip angle = 8°, and receiver bandwidth = 620.8 kHz. For VBM analysis, we used the optimized procedure proposed by (Good et al., 2001). In summary, anatomical images were first segmented into different tissue partitions (J. Ashburner & Friston, 2005). Gray and white-matter images were nonlinearly registered to a MNI template using diffeomorphic image registration algorithm (DARTEL) (J. Ashburner, 2007). To accommodate for brain size differences, registrations were performed iteratively in a coarse-to-fine manner. Volumetric changes of each voxel were obtained by multiplying (or modulating) voxel values in the gray matter image by the deformation field derived from the registration procedure. Individual, modulated images were resampled to 1.5 mm isotropic voxels and spatially smoothed with a 6-mm FWHM kernel. To account for the dependence of participants’ multiple scans in this study, GMV images were analyzed using Sandwich Estimator method, which was designed for analyzing longitudinal and repeated measures data (Guillaume, Hua, Thompson, Waldorp, & Nichols, 2014). The model included group (pCWS and controls) and group by age interaction as well as quadratic age, sex, IQ, brain size, socioeconomic status and stuttering severity as covariates to control potential sources of variation. Although there was a significant difference between CWS and controls in IQ and both language measures, PPVT and EVT (Table 1), only IQ was included in the model because both measures were highly correlated with IQ (PPVT-IQ *r*=0.70, EVT-IQ *r*=0.69). The overall means of each covariate, except stuttering severity, were removed to capture the variation and potential differences between groups associated with the covariates. Since stuttering severity is only relevant to CWS, we considered stuttering severity was zero for controls and the mean for CWS was removed from the measure so that it would remove the variation associated with stuttering severity without affecting the group estimates. Voxel-wise *t*-statistics of the group difference were calculated. For comparing our VBM results with the findings in the literature, we also apply a threshold to visualize the significant GMV differences.

However, it is not the primary goal of this study. Voxel-wise height threshold *p*<0.005 and cluster-size threshold *k*>316 voxels were used to control for false positives. This set of threshold corresponds to a corrected *p*<0.05. The cluster-size threshold was determined by AFNI 3dClustSim (version 17.2.13). Specifically, we first generated a non-gaussian noise model according to the spatial smoothness of the residual images using the AFNI 3dFWHMx autocorrelation function (-acf option). Then, we used Monte Carlo simulations implemented in AFNI 3dClustSim to estimate the false positive rate from the noise model (Cox, Chen, Glen, Reynolds, & Taylor, 2017).

### Gray matter volume and gene expression correlation

Microarray-based gene expression data were obtained from the AIBS, which provides normalized expression of 29,131 genes using a total of 58,692 probes in each of 3,702 brain samples obtained from six adult donors (5 males, 1 female; age: 24-57 years; see http://www.brain-map.org/ for details) (Hawrylycz et al., 2012). We excluded genes whose symbols could not be identified in the database of HUGO Gene Nomenclature Committee, resulting in a total of 19,174 unique genes. The T1-weighted magnetic resonance images of the donors were segmented into different tissue partitions and normalized to the MNI template using the same procedure used for analyzing the structural images acquired from our pediatric subjects. Using the deformation field generated by DARTEL, the locations of brain samples in native space were transformed into the MNI space. The samples’ locations were mapped to 90 cortical and subcortical regions and the cerebellum based on a standard atlas with automated anatomical labeling (AAL Atlas) (Tzourio-Mazoyer et al., 2002). Because samples in the right hemispheres were taken from only two of the six donors, only supratentorial regions in the left hemisphere were included in our GMV-gene expression analysis. Since the right cerebellum has strong anatomical connections with the left cerebral hemisphere, and cerebellar anomalies have been associated with stuttering, the right cerebellum was included in our analyses as a single region. In total, 46 regions were included in the primary GMV-gene expression analysis. For each donor, expression of the same gene at each sample location from different probes was first averaged. Gene expression for each region was represented by the median of all the samples in the region. This step generated a parcellated expression map for each of the 19,174 genes. The GMV difference of each region was calculated by taking the mean of voxel-wise absolute *t*-statistics of between-group GMV differences (|*t*-stat|) within the region. Absolute GMV difference was used because the effect of genetic mutations on GMV is not known. Many neurological disorders such as ADHD and childhood onset schizophrenia are associated with the reduction of GMV (Gogtay et al., 2004; Nakao, Radua, Rubia, & Mataix-Cols, 2011). Like these neurological disorders, the effect of genetics in stuttering could directly impact the function of a brain region, leading to the reduction of GMV in stuttering. Second, the effect of genetics may also delay the cortical developmental trajectories of gray matter in children who stutter. The normal developmental trajectories of GMV differ across different brain areas, with most areas showing general decreases with age (Ducharme et al., 2015; Lange, 2012). Thus, delays in GMV decreases during development may appear as increased GMV when compared with age-matched controls. We do not know how the effect of genetics will lead to decreases or increases of GMV in stuttering, and how the mechanisms underlying the increases and decreases interplay in different brain regions. Moreover, neurological disorders with a strong genetic underpinning can be associated with increased or decreased GMV in different brain regions. For example, autism spectrum disorder (ASD) is associated with a GMV decrease in the bilateral amygdala-hippocampus complex and an increase in the middle-inferior frontal gyrus (Via, Radua, Cardoner, Happé, & Mataix-Cols, 2011). Moreover, a small previous study on GMV in pCWS shows that stuttering is associated with GMV decreases in the inferior frontal gyrus and the putamen as well as increases in other speech motor areas (Beal, Gracco, Brettschneider, Kroll, & De Nil, 2013). Therefore, the effect of genetic variations on GMV could be in different directions, and we chose to use the absolute GMV differences. To minimize the potential adverse effect of outliers, Spearman’s rank correlation was used to assess the relationship between gene expression and between-group GMV difference instead of Pearson’s correlation as it was used in previous analyses of this kind (Ortiz-Terán et al., 2017; Richiardi et al., 2015). Spearman’s correlation coefficients (*ρ*) between GMV differences and each of the 19,174 genes expressed across the 46 regions were calculated. This procedure established a distribution of correlation coefficients. Statistical threshold was set at *q*<0.05 (adjusted *p*<0.05), corrected for multiple testing by controlling the false discovery rate (FDR) (Benjamini & Hochberg, 1995).

### Gene set enrichment analysis

Since genes other than *GNPTG* and *NAGPA* that are expressed in concordance with between-group GMV differences might also be associated with persistent stuttering, we carried out a gene set enrichment analysis to identify biological processes, molecular functions, cellular components or KEGG (Kyoto Encyclopedia of Genes and Genomes) pathways for which the top 2.5% of the genes that were most positively correlated with GMV differences are enriched. The 19,174 genes were used as the input of the background set for the enrichment analyses. We used PANTHER (http://geneontology.org/) to identify enrichment for biological processes, molecular functions or cellular components and DAVID (david.ncifcrf.gov) for identifying enrichment for KEGG pathways (M. Ashburner et al., 2000; Huang, Sherman, & Lempicki, 2009;

The Gene Ontology Consortium, 2017). The redundancy of the resulting gene ontology terms were removed by using REViGO (Supek, Bošnjak, Škunca, & Šmuc, 2011). Fisher’s Exact test was used to determine statistical significance of enrichment factors. Statistical threshold was set at *q*<0.05, corrected for multiple testing by controlling the FDR (Benjamini & Hochberg, 1995). Although the regional expression of our targeted four genes were only positively correlated with the GMV differences, for exploratory purposes, we carried out the same enrichment analysis using the 2.5% of the genes that were most negatively correlated with GMV differences.

## Results

We used 87 longitudinally-acquired structural scans from 26 pCWS and 140 scans from 44 controls using a well-established neuroimaging technique, voxel-based morphometry (VBM) (J. Ashburner, 2007; Good et al., 2001), to estimate voxel-wise GMV differences across the whole brain. Controlling for sex, age, quadratic age, cranial brain volume, IQ, socioeconomic status and stuttering severity, the longitudinal analysis model showed that in the pCWS group, GMV in the left somatosensory, left anterior prefrontal and the right motor areas was significantly larger than in the control group (Fig. 1A).

**Figure 1.**
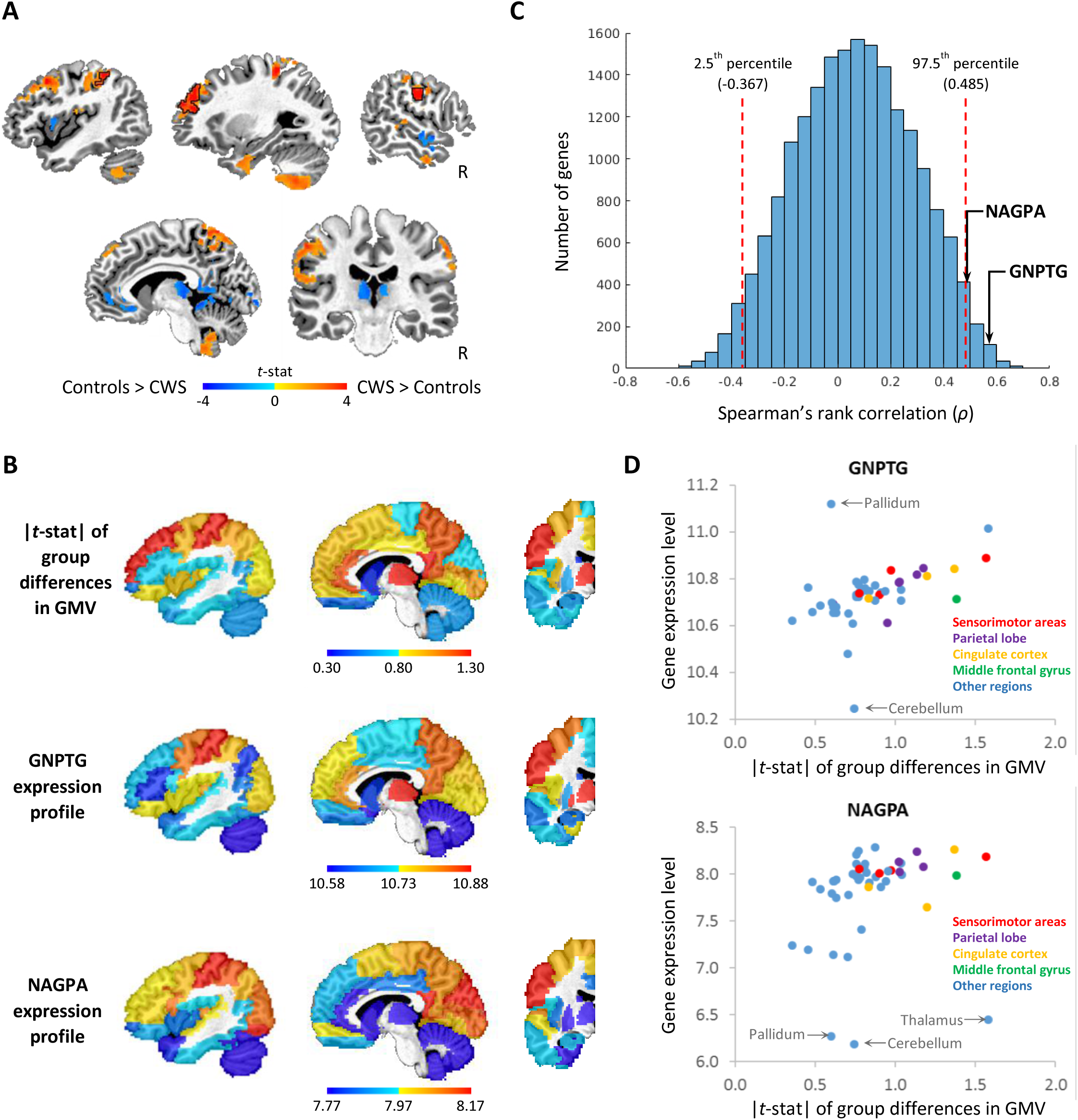
Spatial relationship between gene expression and between-groups differences in gray matter volume (GMV). (A) Voxel-wise differences between children with persistent stuttering (pCWS) and controls in GMV. Color-coded *t* values of group differences are overlaid on an anatomical image. Areas exhibited a significant between-group difference at multiple comparisons corrected *p*<0.05 are outlined by black lines. The other colored areas are subthreshold (uncorrected *p*<0.1). (B) Parcellated gene expression of *GNPTG* and *NAGPA* and absolute GMV differences in *t*-statistics (|*t*-stat|) in 45 left hemispheric regions and the right cerebellum (which anatomically connects to the left hemisphere) were overlaid on a single-subject anatomical image. The parcellation of the brain was based on a standard atlas with automated anatomical labeling (AAL). Gene expression and |*t*-stat| of the right cerebellum were displayed in the left cerebellum to save space. (C) Frequency plot of Spearman’s correlation coefficients between GMV group differences and each of the 19,174 genes expressed across the regions. The red dash lines indicate the levels of correlation at the 2.5^th^ and 97.5^th^ percentiles. To obtain a correlation >0.474 or <-0.360 is less than 5% chance if a gene is randomly selected. (D) Scatter plots between gene expression and group differences in GMV in the sensorimotor areas (red dots), the parietal lobe (purple dots), the cingulate cortex (orange dots), the middle frontal gyrus (green dots) and the rest of the regions (blue dots). Regions in which gene expression is 2.5 standard deviations above or below mean were labelled.

We examined the spatial correspondence between the expression of the *GNPTG, NAGPA, GNPTAB* and *AP4E1* genes and between-group differences in GMV across the 46 regions using the Spearman rank correlation (Fig. 1B). The correlation coefficients (*ρ*) associated with *GNPTG, NAGPA, GNPTAB* and *AP4E1* genes were 0.57 (*q*<0.01), 0.49 (*q*<0.05), −0.08 (*q*=0.78) and 0.08 (*q*=0.78), respectively. As illustrated in the frequency distribution of *ρ* of all 19,174 genes in our analysis (Fig. 1C), the *ρ* associated with *GNPTG* and *NAGPA* genes were significantly higher than the 97.5 percentile. Since there was a tendency for a between-group difference in sex ratio [χ^2^(1, *N*=70)=3.09, *p*=0.079], we conducted a follow up analysis including only male pCWS and controls to rule out the possibility that the observed gene-brain relationship was driven by potential sex differences. The results of the male-only analysis were very similar to the original results (i.e., *GNPTG: ρ*=0.47, *q*<0.05; *NAGPA: ρ*=0.57, *q*<0.01; *GNPTAB*: *ρ*=0.03, *q*=0.93 and *AP4E1 ρ*=0.01, *q*=0.98), indicating that the results were not driven by a higher proportion of female subjects in the control group. The scatter plots in Fig. 1D show the relationship between gene expression and between-group differences in GMV across regions in the original analysis. We observed that the expression level of the *GNPTG* and *NAGPA* genes in the cerebellum and some subcortical regions such as the pallidum deviated from the relationship seen for the other regions. Repeating the analysis excluding the cerebellum, basal ganglia regions and thalamus, we obtained similar results (i.e., *GNPTG*: *ρ*=0.57, *q*<0.05; *NAGPA*: *ρ*=0.42, *q*<0.10; *GNPTAB*: *ρ*=0.02, *q*=0.58 and *AP4E1*: *ρ*=0.08, *q*=0.50), although the correlation for *NAGPA* only showed a tendency toward significance (Fig. S1 in the Appendix). Fig. 1D further illustrates that, in our original analysis, the monotonic relationship between the expression of *GNPTG* and *NAGPA* and between-group differences in GMV in different regions. To further explore whether the relationship between gene expression and between-group differences in GMV was specific to persistent stuttering, we performed the same analysis on 17 children who had recovered from stuttering (rCWS). For rCWS, the relationship was not significant for any of the four genes (i.e., *GNPTG*: *ρ*=-0.08, *q*=0.56; *NAGPA*: *ρ*=0.20, *q*=0.38: *GNPTAB*: *ρ*=-0.24, *q*=0.34; *AP4E1*: *ρ*=0.15, *q*=0.45).

Gene set enrichment analysis of the top 2.5% of the genes whose expression was most positively correlated with the between-group differences in GMV (Table S1 in the Appendix) showed that this set of genes was highly enriched with genes involved in energy metabolism and mitochondrial functions (Table S2-S3 in the Appendix). Genes in several KEGG pathways were also over-represented in the gene set. Forty-nine out of 479 (10.2%) genes analyzed were involved in metabolic pathways, including over representation of a subset of genes involved in citrate cycle (has 00020) and oxidative phosphorylation (hsa00190) (Table S4 in the Appendix). Genes involved in oxidative phosphorylation were also linked to a number of neurological disorders, including Parkinson’s disease, Alzheimer’s disease, Huntington’s disease and amyotrophic lateral sclerosis (Table S5 in the Appendix). Using the same method, the top 2.5% genes negatively correlated with GMV differences were significantly enriched for the DNA packaging complex and the nucleosome (cellular components) as well as three KEGG pathways: Alcoholism (hsa05034), Systemic lupus erythematosus (hsa05322) and Viral carcinogenesis (has05203) (Table S6 and S7 in the Appendix).

## Discussion

In our VBM analysis, we observed significant GMV increases in the left somatosensory areas, the left middle frontal gyrus and the right motor cortex in pCWS. Moreover, the magnitude of absolute GMV differences across brain regions in pCWS was positively correlated with the expression levels of two stuttering-related genes (*GNPTG* and *NAGPA*). This association suggests that mutations in these genes (and potentially related yet to be identified stuttering-related genes) are associated with the changes in GMV in pCWS and the manifestation of persistent stuttering.

To date, two VBM studies in children who stutter have been published in peer reviewed journals (Beal et al., 2013; Chang, Erickson, Ambrose, Hasegawa-Johnson, & Ludlow, 2008). However, the sample sizes of both studies are small (<12 pCWS) and relatively lenient statistic thresholds were used. Nevertheless, both studies showed that smaller GMV in the bilateral inferior frontal gyrus (IFG) is associated with pCWS. In the current study, decreased GMV in the IFG was observed only at an uncorrected threshold (Fig 1A). On the other hand, our data showed that pCWS were associated with significantly increased GMV in the left prefrontal and bilateral sensorimotor areas, which have been shown to have increased blood flow during speech production in people who stutter in previous PET studies (Braun et al., 1997; Fox et al., 1996). This finding is partially consistent with Beal et al. (2013), where pCWS exhibited increased GMV in right motor regions. From previous VBM studies in typically developing children, it is known that a majority of brain regions undergo GMV decreases during childhood, which reflects refinements of neural circuits via synaptic and dendritic pruning (Gennatas et al., 2017). The increased GMV in pCWS relative to controls may thus reflect a delay of development in those areas. Similarly, developmental delays in the structure of white matter tracts connecting speech-motor areas have been observed in a previous diffusion tensor imaging study using the same group of subjects (Chow & Chang, 2017). However, the connection between white and gray matter anomalies is unclear and warrant further analysis.

While our results are correlational and do not provide direct evidence on the underlying casual mechanisms, here we discuss biological mechanisms that could explain to the observation of our results. Our analysis with regional gene expression showed that the magnitudes of GMV regional differences between pCWS and controls are correlated with the expression patterns of *GNPTG* and *NAGPA* in the left hemisphere and the right cerebellum. This finding supports our hypothesis that the altered GMV in pCWS is related to known genes associated with stuttering. *GNPTAB*, *GNPTG* and *NAGPA* are involved in the formation of the M6P tag that allows the binding of the lysosomal enzymes and other M6P-glycoproteins to the M6P receptor. While the main function of the M6P receptor is to transport the M6P-hydrolases from the trans Golgi network (TGN) to the lysosomes, M6P receptors also bind other M6P-proteins and non-M6P proteins such as the insulin growth factor 2 (IGF2), a hormone that regulates cell metabolism and growth at the cell surface (Barnes et al., 2011; Gary-Bobo, Nirdé, Jeanjean, Morère, & Garcia, 2007; Han, D’Ercole, & Lund, 1987). The cation-independent M6P receptor plays an important role in the regulation of IGF2 levels by mediating its internalization and degradation (Oka, Kawasaki, & Yamashina, 1985). While IGF2 binds at a different site on the receptor than M6P tagged proteins, M6P lysosomal enzymes have been shown to alter the binding of IGF2 to the receptor (De Leon, Terry, Asmerom, & Nissley, 1996; Kiess et al., 1989). Thus, altered binding of these enzymes due to mutations associated with stuttering might alter IGF2 mediated growth leading to altered GMV in regions where these enzymes are highly expressed. IGF2 has been linked to brain growth and differentiation as well as psychiatric and neurodegenerative disorders such as anxiety disorders and Parkinson’s disease (Fernandez & Torres-Alemán, 2012; Matrone et al., 2016; Pardo et al., 2018). While speculative, the connection between stuttering-related genes and IGF2 would be an interesting future research direction.

The gene set enrichment analysis showed that genes expressed in concordance with the between-group differences in GMV were significantly enriched for metabolic processes in mitochondria (Table S2-4 in the appendix). Moreover, the gene set also enriched for the KEGG pathways of a number of neurological disorders, including Parkinson’s disease, Alzheimer’s disease, Huntington’s disease and amyotrophic lateral sclerosis (Table S5 in the appendix). The genes involved in these disease pathways largely overlapped with genes involved in oxidative phosphorylation, suggesting that metabolic dysfunction similar to these diseases may play a role in stuttering. The positive correlation between GMV differences and expression of metabolic genes indicates that the regions exhibiting large GMV differences have higher energy metabolism rates (Goyal, Hawrylycz, Miller, Snyder, & Raichle, 2014). This result suggested that there may be a link between energy metabolism and the development of anomalous GMV in persistent stuttering. This link could be mediated through mutations in metabolic genes as candidates in persistent stuttering. However, it is also possible that the GMV differences between pCWS and controls were exacerbated in brain regions with relatively high energy consumption related to disturbance in lysosomal function (McKenna, Schuck, & Ferreira, 2018).

How might metabolism and gene mutations that affect lysosomal enzymes trafficking lead to the neurological anomalies associated with stuttering? Recent studies have shown that lysosomes and mitochondria interact physically and functionally, and these interactions play an important role in modulating metabolic functions of the two organelles (Todkar, Ilamathi, & Germain, 2017; Wong, Ysselstein, & Krainc, 2018). The known gene mutations related to stuttering are involved in the trafficking of lysosomal enzymes for the breakdown of macromolecules and organelles, including damaged mitochondria (Plotegher & Duchen, 2017). If the lysosomal function cannot keep pace with the sharp increase of energy metabolism, damaged mitochondria may accumulate, leading to increased oxidative stress, which may in turn negatively impact neurological development (de la Mata et al., 2016; Kiselyov, Yamaguchi, Lyons, & Muallem, 2010). The effect of this vulnerability may be amplified in brain regions with relatively high energy consumption in children between 2 to 5 years of age, a period of time in which brain metabolism sharply increases (Chugani, Phelps, & Mazziotta, 1987; Goyal et al., 2014). This age range also coincides with the typical onset age of stuttering (Bloodstein & Ratner, 1995), as well as a period of rapid development of language and other cognitive functions.

Consistent with this hypothesis, significant increases in mitochondrial fragmentation were observed in several lysosomal storage diseases, including Mucolipidosis Types II and III that are caused by homozygous mutations in the same genes linked to stuttering (*GNPTAB* and *GNPTG*) (Jennings et al., 2006). Although mutations in people who stutter are usually heterozygous and in different locations and forms (C. Kang et al., 2010) and do not lead to the detrimental symptoms of Mucolipidosis, a previous study showed that lysosomal enzyme activity is partially compromised by mutations associated with persistent stuttering in *NAGPA* (Lee, Kang, Drayna, & Kornfeld, 2011). However, it is unclear to what extent cellular function is affected by a partial deficiency of lysosomal enzymes, especially related to mitophagy and autophagy.

While our study showed a significant spatial relationship between expression patterns of *GNPTG*/*NAGPA* and GMV regional differences between pCWS and controls, a few methodological caveats should be acknowledged. First, although the sample size of the current study is one of the largest neuroimaging data sets in developmental stuttering and included repeated scans from each subject, it is small in the context of neuroanatomical studies examining correlates in complex heterogeneous traits. Small sample sizes could lead to finding spurious group differences (Fusar-Poli et al., 2014) as well as lower power that limit detection of subtle differences that might otherwise be possible with larger samples. Second, although gene expression data from AIBS were obtained from six adult donors, there is a high degree of similarity in the regional expression among donors (Hawrylycz et al., 2012). Moreover, the microarray gene expression data provided by AIBS were normalized within and across donors’ brains. The details of the current normalization method can be found in a technical paper from Allen Brain Atlas (http://help.brain-map.org/download/attachments/2818165/Normalization_WhitePaper.pdf). This normalization procedure reduces the variability across donors due to technical biases and allows comparison of expression data across two or more brains. However, a certain degree of variability between donors is inevitable. To further explore the individual donor variability, we examined the spatial relationship between the GMV differences with the express of the four gene in each of the six donors. The correlation coefficients are presented in (Table S8 in the Appendix). In summary, we found that the correlation with *GNPTG* and *NAGPA* was positive for all six donors and at least one of them in each donor was larger than 0.30 (*p*<0.05) in four of the six donors, whereas the correlation with *AP4E1* and *GNPTAB* ranged from −0.23 to 0.20 (*p*>01). While variability of individual donors in gene expression is inevitable, the results of this individual donor analysis point to the same direction of our results using the gene expression aggregated from the six donors. For readers who are interested in further examining the individual donor variability, the regional expression patterns of the four targeted genes in each of the six donors are presented in Fig. S2 and S3 in the Appendix. To date, the expression data from AIBS is the only source of human gene expression patterns with high spatial resolution. As mentioned in the introduction, AIBS gene expression data and the approach of the current study have been used to reveal relationships between genes and intrinsic brain network, association between neuroplasticity-related genes and altered functional connectivity in blind children, and to confirm the relationship between known risk genes and their associated neurological disorders including Alzheimer’s disease, Huntington’s disease and schizophrenia (Grothe et al., 2018; McColgan et al., 2017; Ortiz-Terán et al., 2017; Richiardi et al., 2015; Romme et al., 2017). Third, we cannot completely rule out the possibility that expression levels and patterns of some genes in children and adults are different. If this is the case for the genes associated with stuttering, we would expect that the spatial correlation between gene expression in adults and the GMV patterns in children would be weak, whereas a strong relationship was observed in our study. Moreover, previous studies have suggested that the changes of gene expression occur predominately during prenatal and infant development and become relatively stable by around 6 years of age (H. J. Kang et al., 2011). Kang et al. (2011) estimated that only 9.1% of genes exhibit temporally differential expression in the first 20 years of life, and the portion of differentially expressed genes should be even less in our participant’s age group. Future studies using gene expression profiles in children should be pursued to refine our understanding of the relationship between brain anomalies and gene expression, when such dataset becomes available.

## Conclusions

In conclusion, we showed that relative to controls, pCWS exhibited larger GMV in the left somatosensory, left anterior prefrontal and the right motor areas. The spatial pattern of this GMV difference was positively correlated with the expression of lysosomal targeting genes *GNPTG* and *NAGPA* as well as genes involved in energy metabolism. More research is warranted to further investigate possible roles of M6P mediated intracellular and extracellular trafficking as well as metabolic functions play a role in the development of brain structural anomalies associated with stuttering.

## Acknowledgements

The authors wish to thank all the children and parents who participated in this study. We also thank Kristin Hicks for her assistance in participant recruitment, behavioral testing, and help with MRI data collection, Scarlett Doyle for her assistance in MRI data acquisition, and Ashley Diener for her assistance in speech data analyses.

## Appendix

**Fig. S1.**
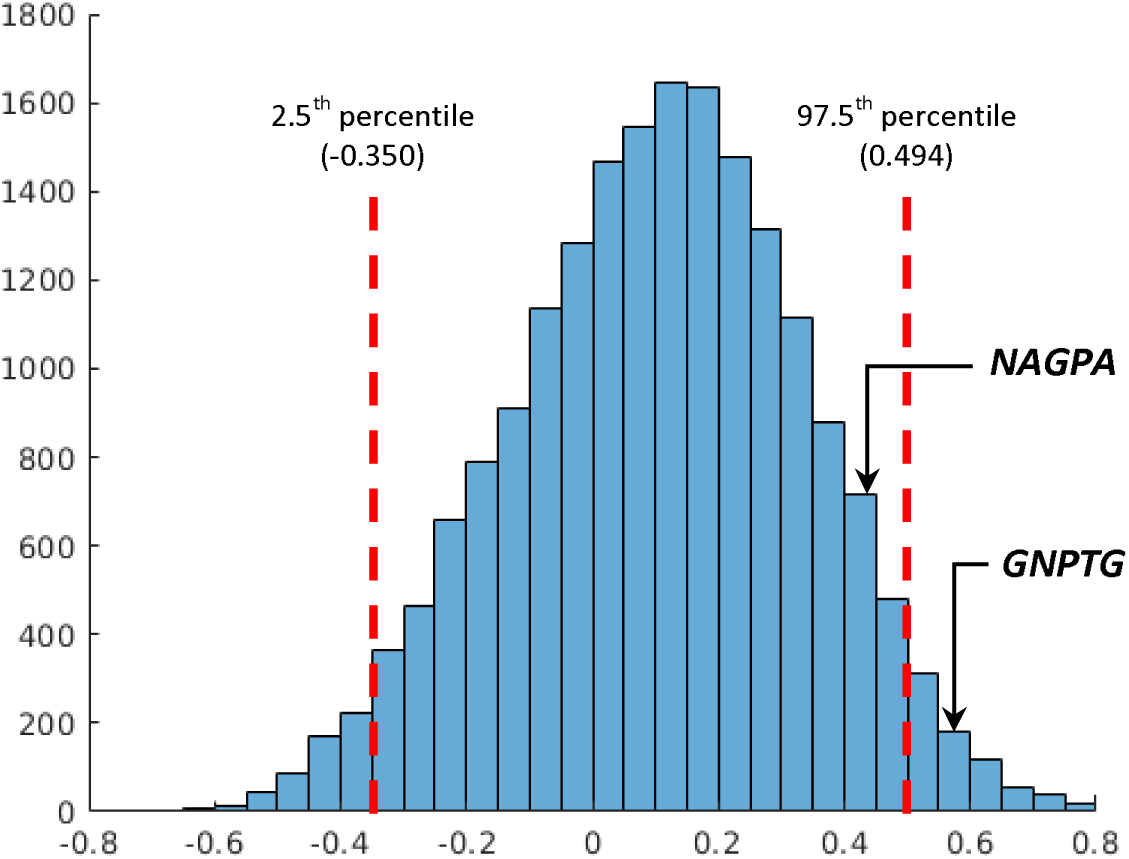
Frequency plot of Spearman’s correlation coefficients between GMV group differences and each of the 19,174 genes expressed across the regions when cerebellum, basal ganglia regions and thalamus were excluded. The red dash lines indicate the levels of correlation at the 2.5th and 97.5th percentiles. To obtain a correlation >0.494 or <-0.350 is less than 5% chance if a gene is randomly selected.

**Fig. S2.**
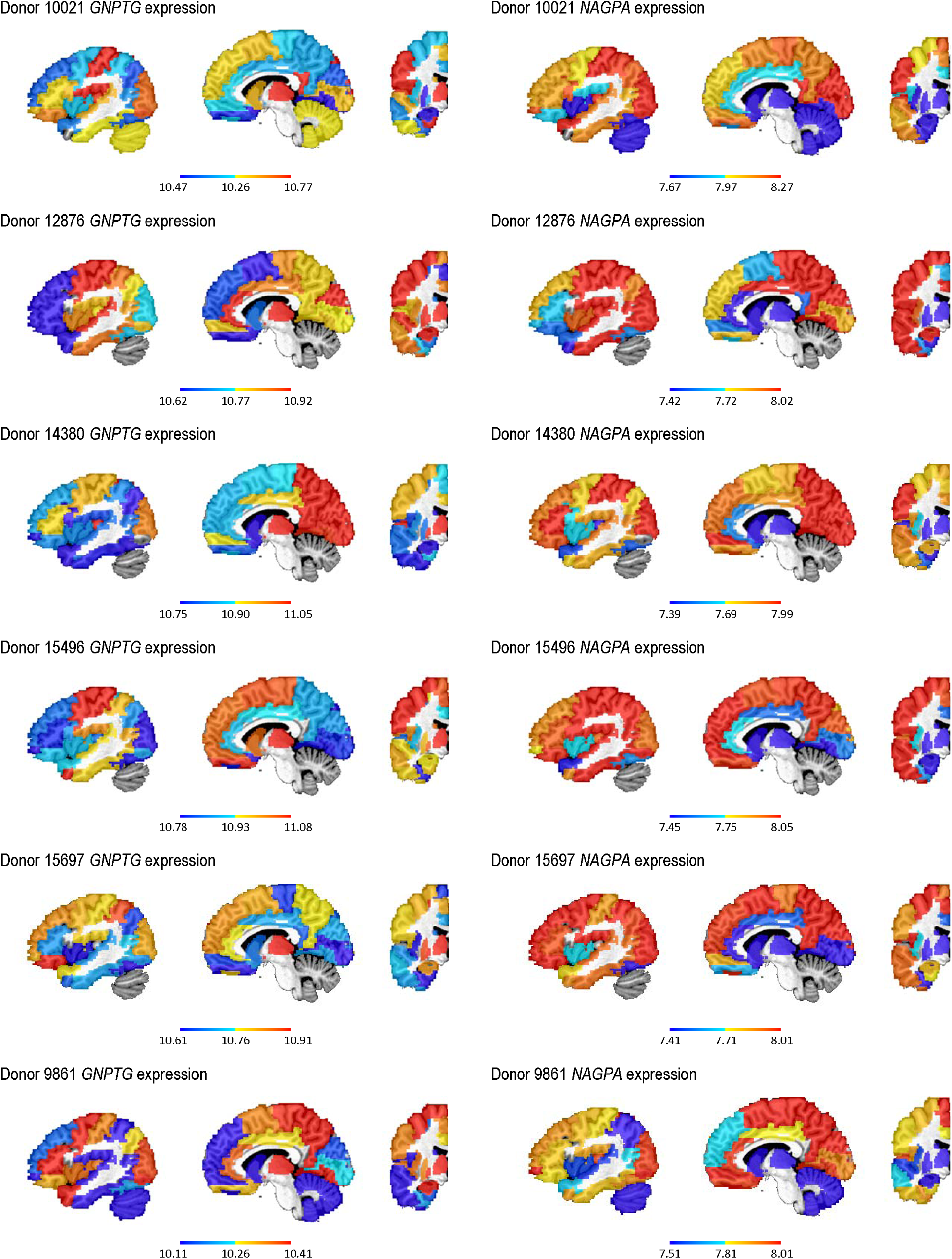
Expression of the GNPTG and NAGPA in each of the six donors

**Fig. S3.**
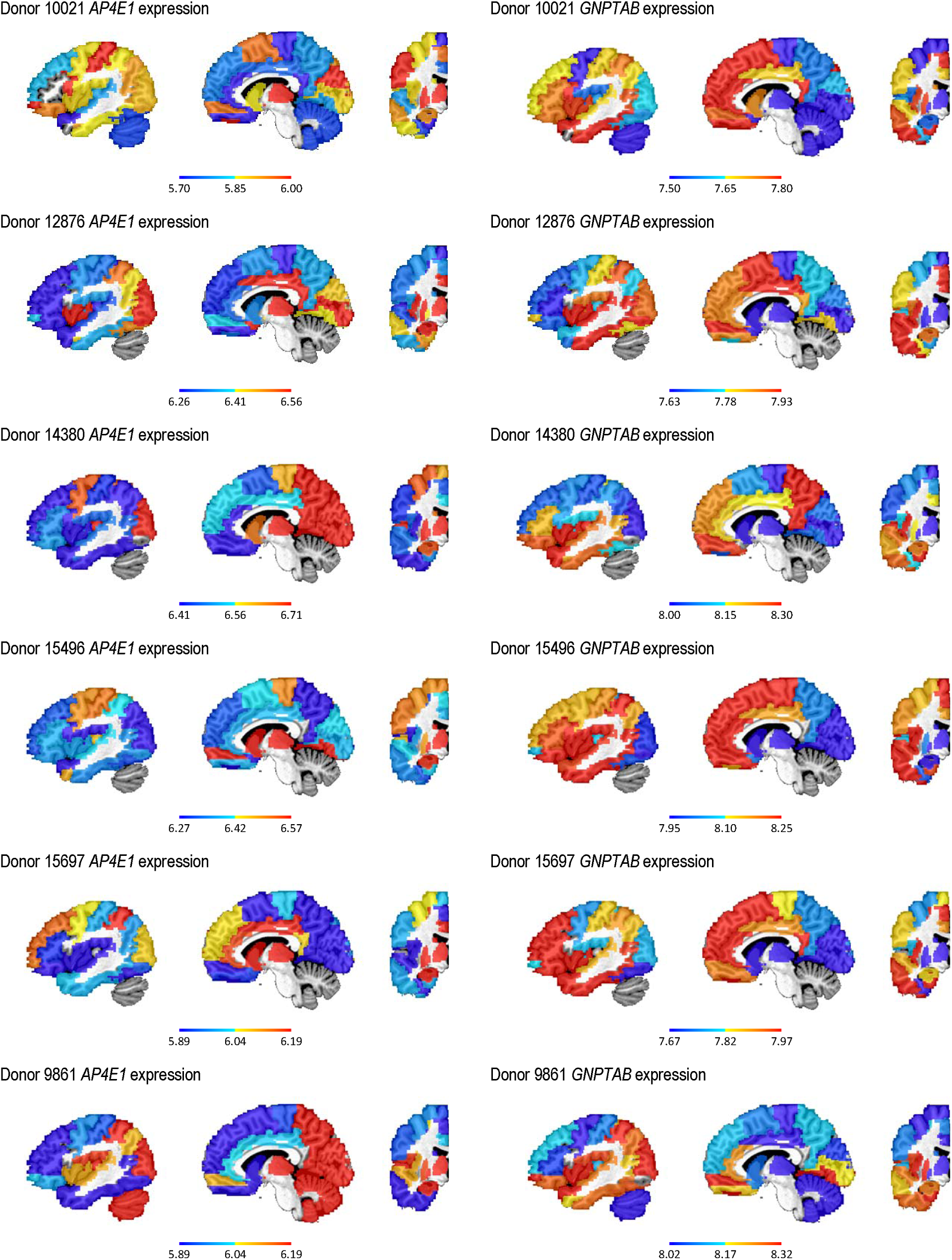
Expression of the AP4E1 and GNPTAB in each of the six donors

**Table S1.**
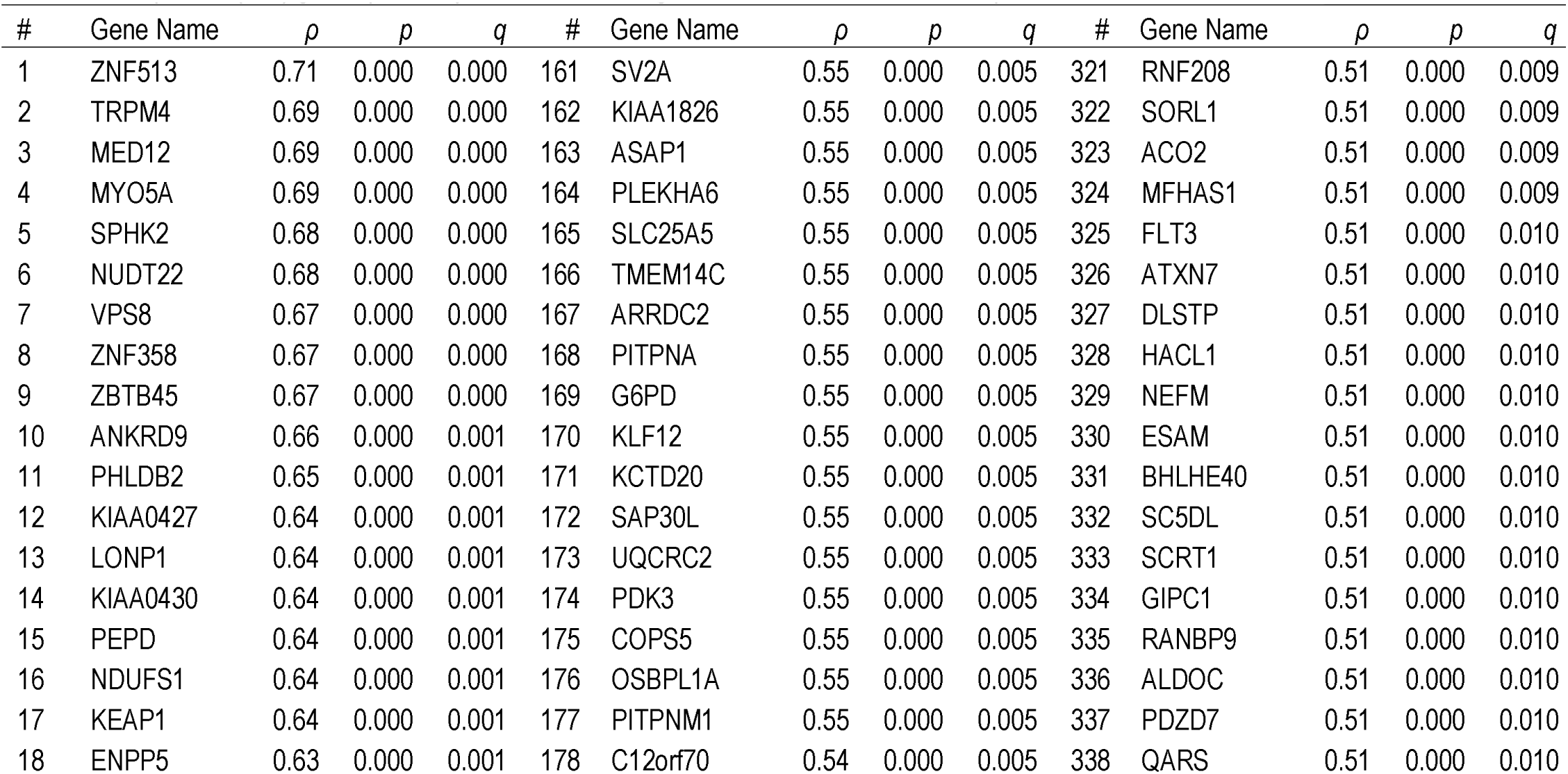
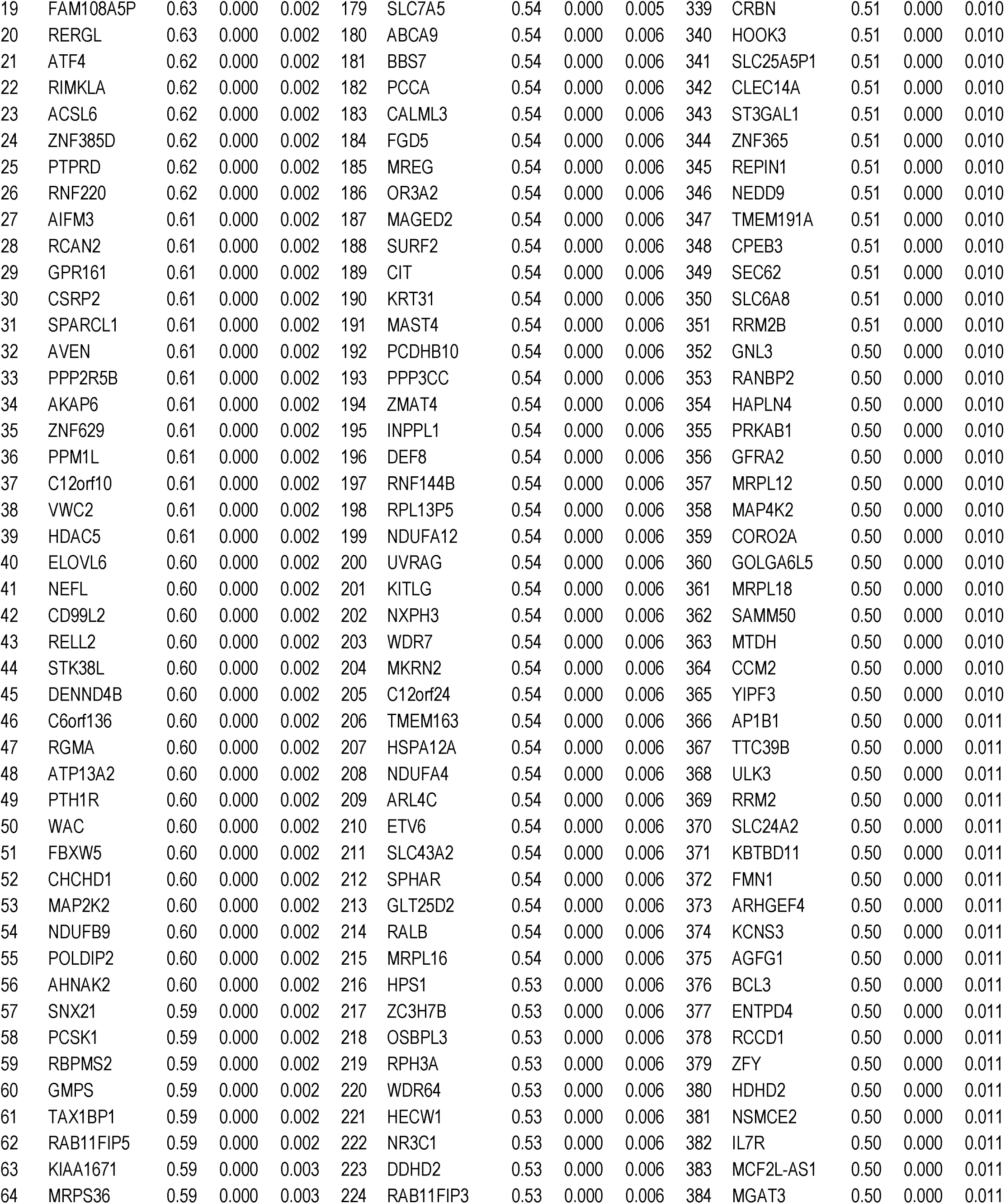
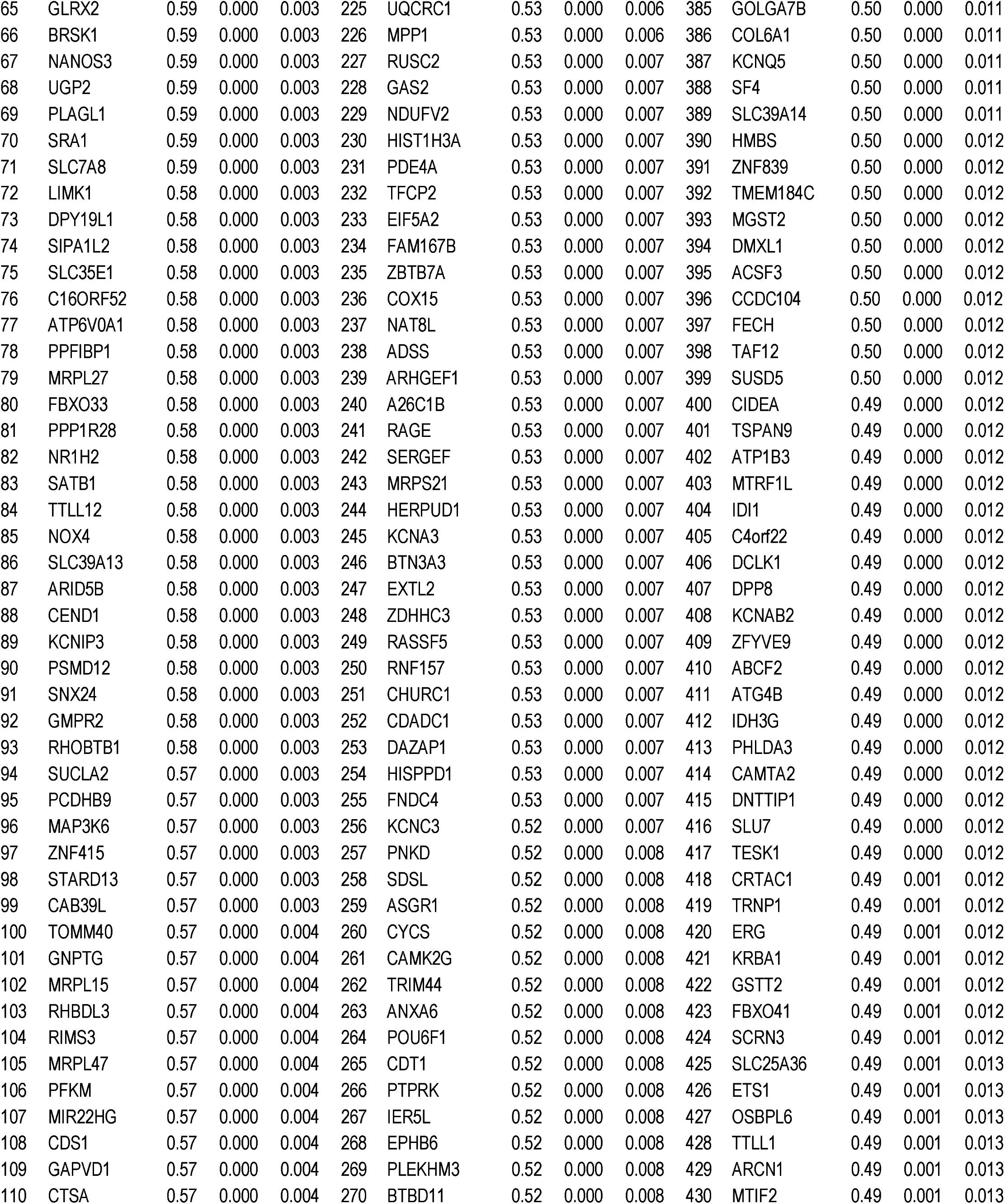
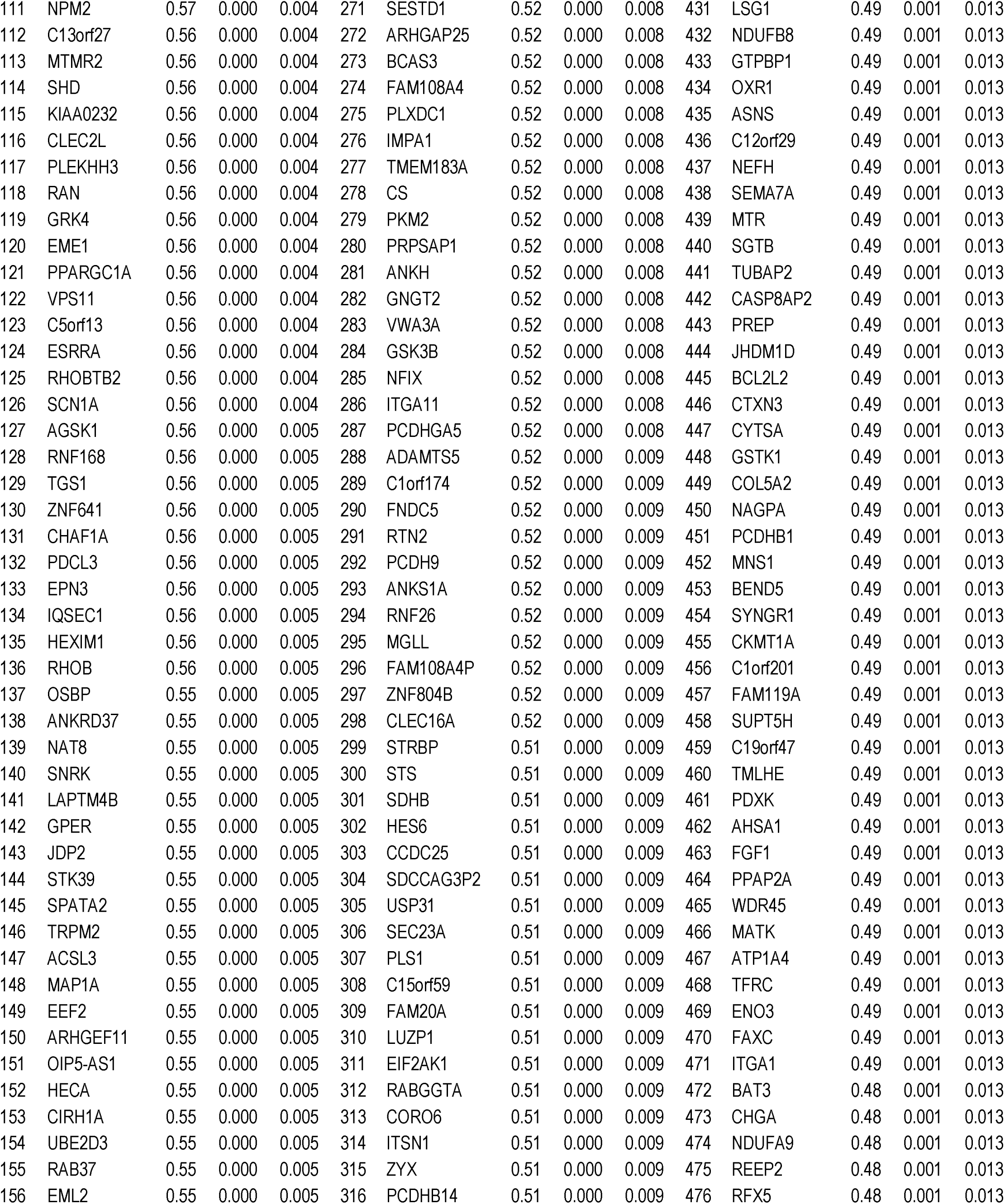
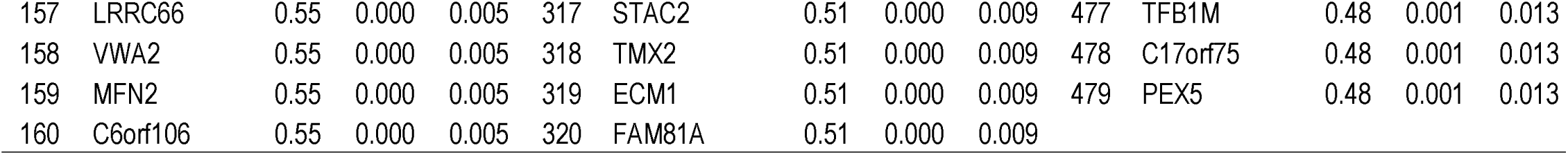
Top 2.5% (479) genes positively correlated with regional GMV difference between pCWS and controls

**Table S2.** Gene ontology (GO) terms for biological processes associated with in the top 2.5% of the genes whose expression was positively correlated with the between group gray matter volume differences.

| GO Term <sup>a</sup> | GO biological process | p Value | FDR q Value | Enrichment <sup>b</sup> | B | b |
| --- | --- | --- | --- | --- | --- | --- |
| GO:0045333 | cellular respiration | 1.02E-06 | 1.62E-02 | 4.43 | 147 | 17 |
| GO:0006414 | translational elongation | 8.37E-06 | 2.21E-02 | 4.46 | 120 | 14 |
| GO:0006091 | generation of precursor metabolites and energy | 2.31E-05 | 3.05E-02 | 2.57 | 387 | 26 |
| GO:0032543 | mitochondrial translation | 1.69E-05 | 2.67E-02 | 4.48 | 111 | 13 |
| GO:0006415 | translational termination | 2.17E-05 | 3.11E-02 | 4.73 | 97 | 12 |

<sup>a</sup> GO terms are arranged hierarchically with the most specific subclass first. GO terms associated with more than 1,000 genes are excluded.
<sup>b</sup> Enrichment = [Number of genes associated with a specific GO term in the target list (b) / Total number of genes in the target list] / [Total number of genes associated with a specific GO term (B) / Total number of background genes].

**Table S3.**
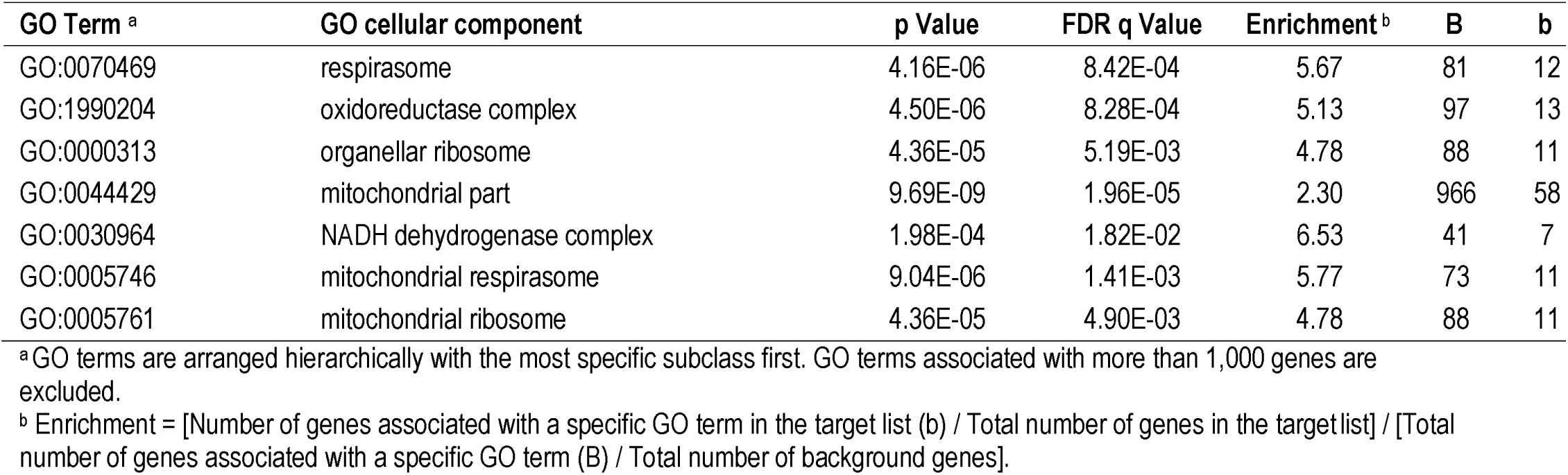
Gene ontology (GO) terms for cellular components associated with the top 2.5% of the genes whose expression was positively correlated with the between group gray matter volume differences.

**Table S4.** KEGG Pathways associated with the top 2.5% of the genes whose expression was positively correlated with the between group gray matter volume differences.

| KEGG ID | KEGG Pathway | p Value | FDR q Value | Enrichment <sup>a</sup> | B | b |
| --- | --- | --- | --- | --- | --- | --- |
| hsa01200 | Carbon metabolism | 4.10E-05 | 1.72E-03 | 4.3 | 112 | 13 |
| hsa05012 | Parkinson's disease | 5.30E-04 | 3.82E-03 | 3.5 | 126 | 12 |
| hsa04932 | Non-alcoholic fatty liver disease (NAFLD) | 1.40E-04 | 5.03E-03 | 3.6 | 146 | 14 |
| hsa05010 | Alzheimer's disease | 3.60E-04 | 3.82E-03 | 3.2 | 161 | 14 |
| hsa01230 | Biosynthesis of amino acids | 5.20E-04 | 3.55E-03 | 4.8 | 70 | 9 |
| hsa00190 | Oxidative phosphorylation | 3.20E-04 | 4.68E-03 | 3.7 | 119 | 12 |
| hsa00020 | Citrate cycle (TCA cycle) | 1.10E-03 | 3.82E-03 | 7.4 | 30 | 6 |
| hsa05016 | Huntington's disease | 3.60E-03 | 2.48E-02 | 2.6 | 183 | 13 |
| hsa00480 | Glutathione metabolism | 1.00E-02 | 3.65E-02 | 4.4 | 50 | 6 |
| hsa05014 | Amyotrophic lateral sclerosis (ALS) | 1.00E-02 | 3.34E-02 | 4.4 | 50 | 6 |

<sup>a</sup> Enrichment = [Number of genes associated with a specific KEGG pathway in the target gene set (b) / Total number of genes in the target set] / [Total number of genes associated with a specific KEGG pathway (B) / Total number of background genes]. KEGG; Kyoto Encyclopedia of Genes and Genomes

**Table S5.**
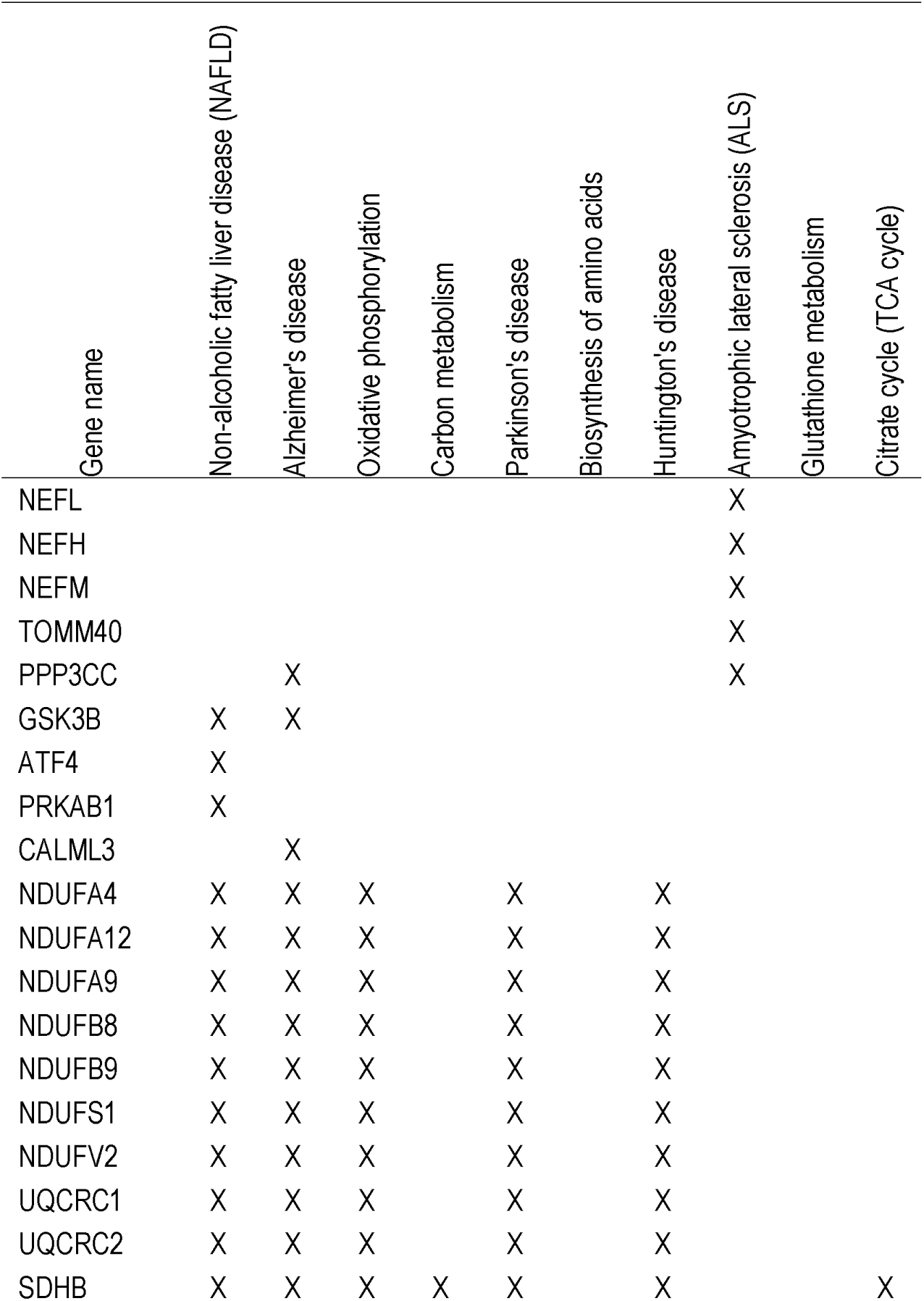
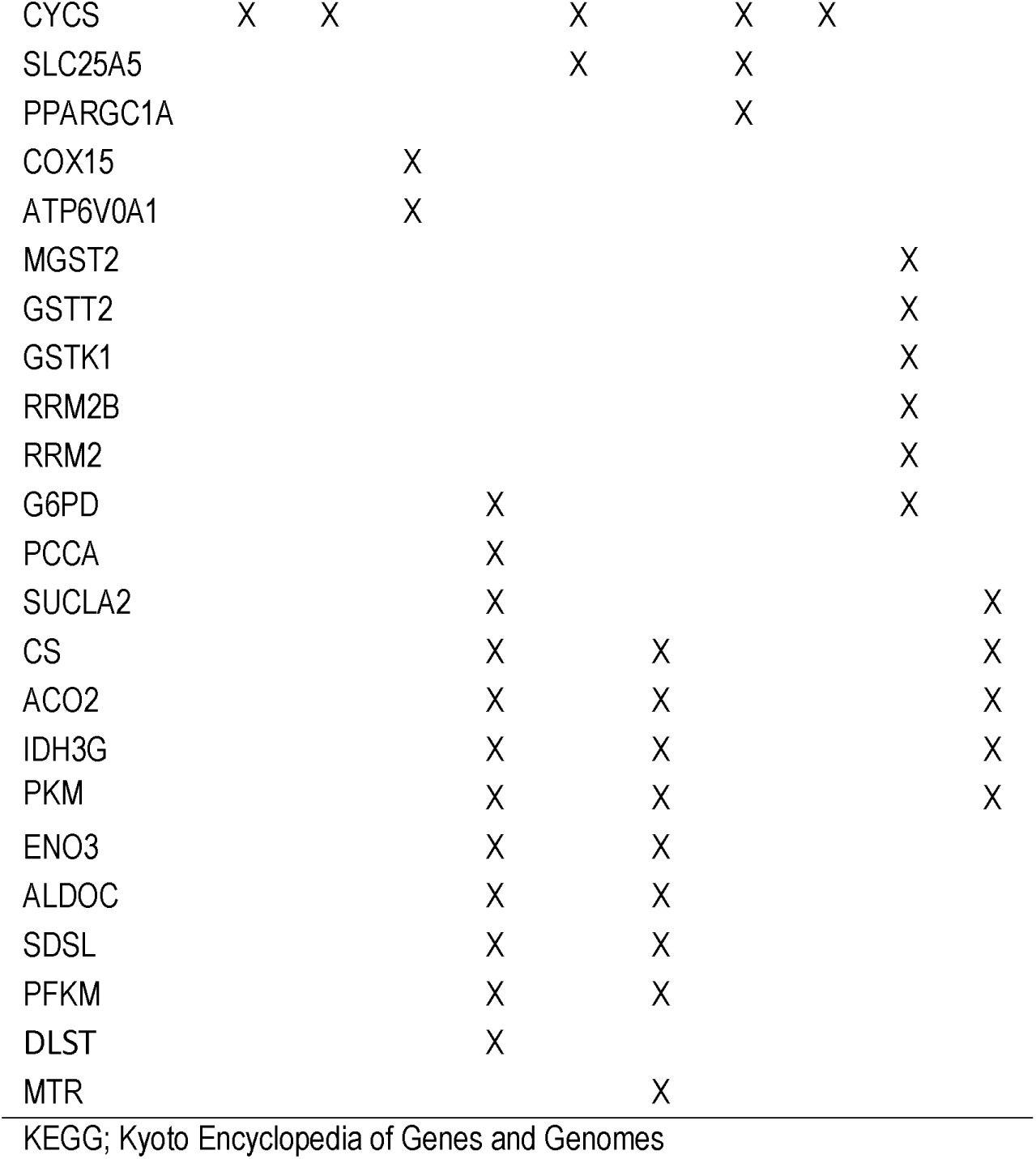
Genes associated with KEGG pathways and between group gray matter volume differences. The statistics of the enrichment analysis that identified the KEGG pathways is listed in Table S4.

**Table S6.**
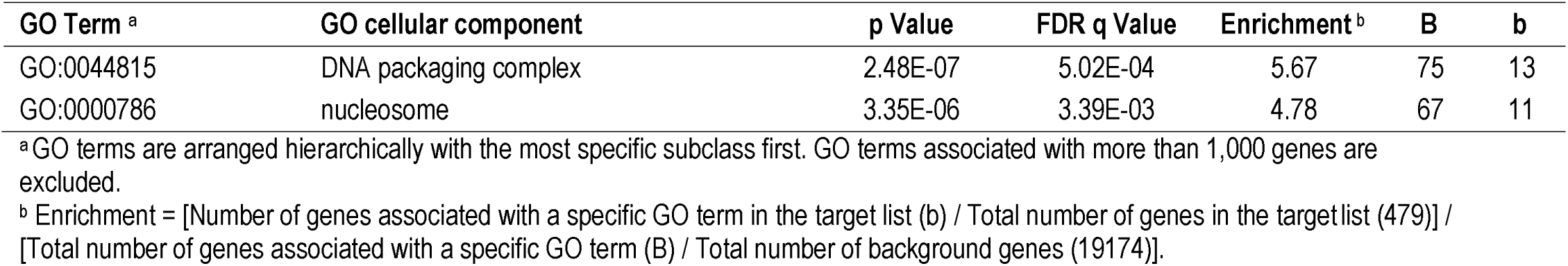
Gene ontology (GO) terms for cellular components associated with the top 2.5% of the genes whose expression was negatively correlated with the between group gray matter volume differences.

**Table S7.** KEGG Pathways associated with the top 2.5% of the genes whose expression was negatively correlated with the between group gray matter volume differences.

| KEGG ID | KEGG Pathway | p Value | FDR q Value | Enrichment <sup>a</sup> | B | b |
| --- | --- | --- | --- | --- | --- | --- |
| hsa05034 | Alcoholism | 4.30E-07 | 9.96E-06 | 4.7 | 170 | 17 |
| hsa05322 | Systemic lupus erythematosus | 1.10E-05 | 1.26E-04 | 4.9 | 126 | 13 |
| hsa05203 | Viral carcinogenesis | 3.10E-03 | 4.00E-02 | 2.8 | 200 | 12 |

<sup>a</sup> Enrichment = [Number of genes associated with a specific KEGG pathway in the target gene set (b) / Total number of genes in the target set] / [Total number of genes associated with a specific KEGG pathway (B) / Total number of background genes]. KEGG; Kyoto Encyclopedia of Genes and Genomes

**Table S8.** Spearman’s correlation between regional GMV differences and the expression of the four targeted genes in the six donors.

| Donor ID | *GNPTG* | *NAGPA* | *AP4E1* | *GNPTAB* |
| --- | --- | --- | --- | --- |
| 9861 | 0.00 | 0.12 | 0.20 | -0.23 |
| 10021 | 0.36 | 0.32 | 0.03 | 0.03 |
| 12876 | 0.18 | 0.23 | -0.01 | -0.02 |
| 14380 | 0.48 | 0.37 | 0.02 | -0.18 |
| 15496 | 0.02 | 0.33 | -0.08 | 0.03 |
| 15697 | 0.15 | 0.32 | -0.02 | 0.11 |

